# An improved framework for grey-box identification of biological processes

**DOI:** 10.1101/2025.02.14.638382

**Authors:** Prem Jagadeesan, Karthik Raman, Arun K Tangirala

## Abstract

Developing models from observations is at the heart of empirical science. Grey-box Modeling combines the insights gained from the results obtained from first principles with observational data. When the model turns out to be unsatisfactory, the goodness of such grey-box models in terms of predictability and parameter estimates largely depends on either modifying the model structure obtained from the first principles or conducting new experiments. Unfortunately, in the context of biological models, where the model structures are usually nonlinear ODEs with a large number of states and parameters along with sparse and noisy experimental data, traditional identification protocols have to go through several iterations to identify the source of the issue. Even after multiple iterations, they may still arrive at sub-optimal solutions. In this work, we propose an improved framework with a new set of tools to resolve this issue unambiguously with a minimum round of iterations.

## 1 Introduction

Data-driven process Modeling (System Identification) has a wide range of applications in medicine and life sciences, such as in pharmacometrics or Quantitative Systems Pharmacology (QSP) [1], discovering drug targets [2], and uncovering the underlying mechanisms of a biological process [3, 4]. In this work, we focus on the grey-box identification of biological processes. A Modeling exercise is considered a grey box if it is possible to incorporate prior knowledge into the modeling exercise in the form of a distribution, a range of parameter values or a model structure dictated by the science of the processes or by order of the dynamical systems etc. However, here we focus on scenarios where the biology of the process dictates the model structure.

System identification is an iterative process and has four major stages: (i) data generation, (ii) model selection, (iii) parameter estimation, and (iv) model validation. To obtain a satisfactory model, tuning may be required in any of these stages depending upon where the problem is and what controllables we have, such as designing new experiments, availability of other candidate models and parameter ranges, etc. The goodness of any model is usually assessed by the goodness of the prediction and the goodness of the parameter estimates [5]. In the case of nonlinear grey-box Modeling, the factors influencing the goodness of the predictions and parameter estimates are not straightforward and there are challenges at each stage of the identification exercise [6, 7, 8]. One of the significant factors affecting the quality of parameter estimates is structural identifiability. A model structure is structurally unidentifiable if two different parameter sets result in identical predictions [5]. Several analytical and computational methods have been proposed to detect the structural identifiability of nonlinear ODEs [9]. Another common factor that affects the goodness of the parameter estimates is sloppiness. Sloppiness is characterized by the presence of large regions in the parameter space that result in nearly identical predictions. The presence of sloppiness will greatly affect the quality of the parameter estimates and sometimes the quality of predictions also [10]. Last but not least, the information contained in the data with respect to a parameter or a model structure impacts the identifiability of the parameters and the goodness of the predictions [5]; lack of information in the data could affect the identifiability of the parameters, sloppiness and the quality of predictions [10, 11]. To overcome this difficulty, generally, Optimal Experiment Design (OED) is proposed, to select experiments that produce informative data [12].

A formal approach to systematically investigate these challenges is proposed in [8, 13, 14]. Given a data set *z* and a model structure (*M*) or set of candidate models (ℳ), the traditional procedure starts with structural identifiability analysis to check whether the given model structure is structurally identifiable, followed by parameter estimation. As a next step, sensitivity analysis is usually done to detect the sensitivity of the parameters at the optimal parameter values. Once the sensitivity analysis is done, the goodness-of-fit to the data is assessed. When the model passes the goodness-of-fit test, the precision of the parameter estimates is analyzed; if both the predictions and parameter estimates are satisfactory, then the model is accepted. Fig. 1 illustrates the flow of the traditional framework. However, this framework must address a crucial problem frequently encountered in grey-box identification. In a Modeling exercise where the model structure is nonlinear, if one encounters poor precision in the parameter estimates as a result of sloppiness analysis, the traditional framework cannot identify the source as it does not include sloppiness analysis prior to estimation. Even in some instances of sloppiness analysis, post estimation, delineating whether the source of the problem lies in the model structure or data becomes impossible [15].

**Figure 1:**
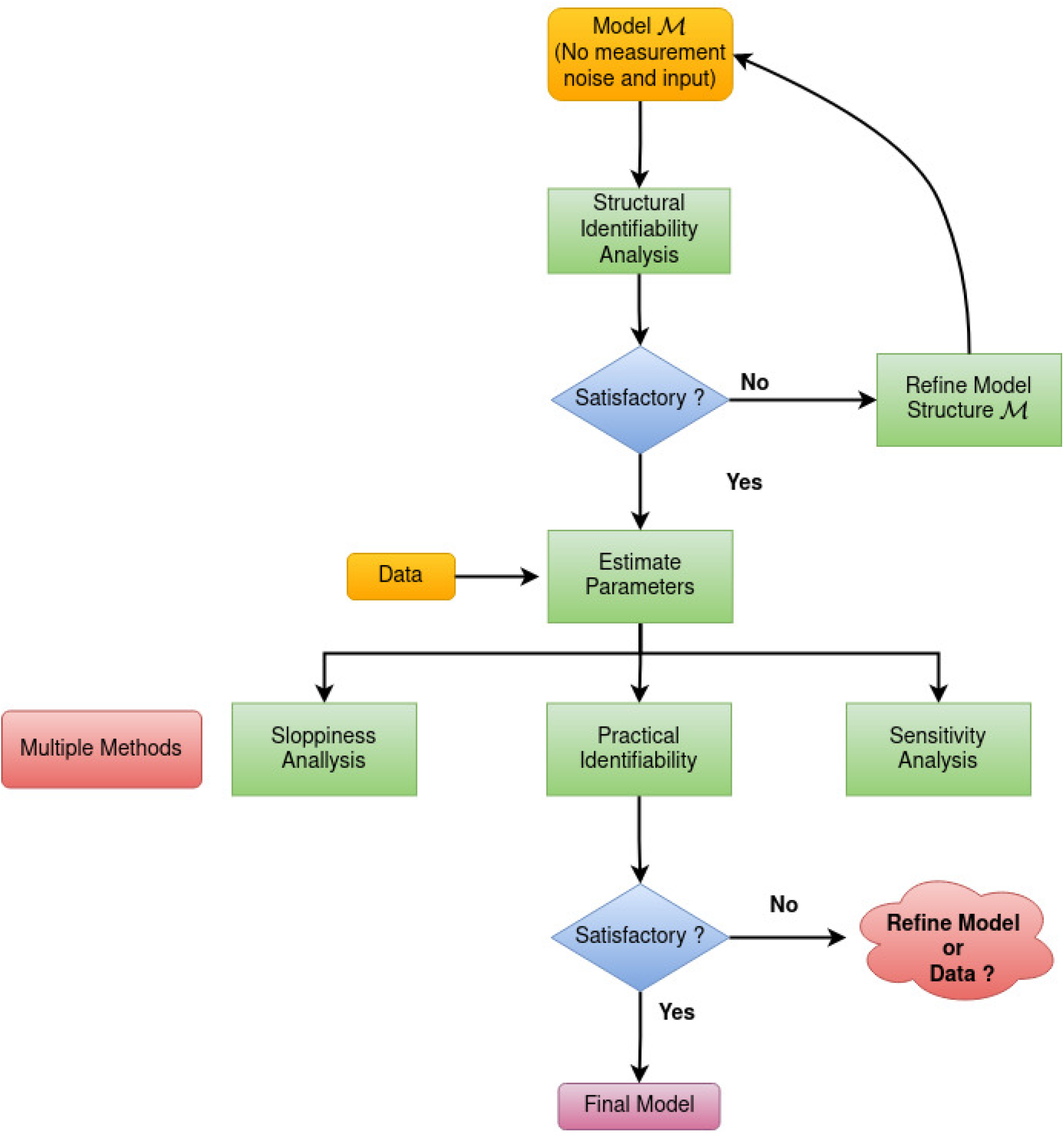
Traditional framework for grey-box identification

In this work, we propose a new framework inspired by our recent method [15] to circumvent the issues discussed above and to identify the source of parameter and prediction uncertainty precisely. The efficiency of the proposed framework is demonstrated using an mRNA network which is multi-scale/sloppy. The proposed method accurately identified the model as the source of parameter and prediction uncertainty. The rest of the paper is organized as follows. Section 3 illustrates the traditional framework and the proposed new framework. Section 4 demonstrates the working of the proposed method using a numerical example, and the paper ends with some concluding remarks in Section 5.

## 2 Methodology

In this section, we discuss the traditional protocol for grey-box identification in detail and the challenges involved in identifying the source of uncertainty. Further, we discuss the new framework in detail that circumvent the issue.

### 2.1 Traditional Framework

The traditional protocol consists of six stages and a corrective mechanism for each stage. Fig 1, shows the workflow that is largely followed in the literature [14, 8, 12]. A brief discussion of each step is given below:

#### Step 1: Structural Identifiability Analysis

Given a model structure and data, the first step is to check for the presence of structurally unidentifiable parameters and their relationships. The parameters can be locally or globally unidentifiable. A brief overview of methods for assessing structural identifiability is given in [9]. Once unidentifiable parameters are detected, one of the following corrective steps can be taken (i) One may choose an alternate model structure or re-parameterize the model (ii) Restrict the parameter search space such that the model is locally identifiable.

#### Step 2: Parameter Estimation

The next step is to estimate the parameters using a particular estimation algorithm suitable for the given problem. Due to the complexity of the cost surface, the choice of optimization algorithm also influences the quality of the parameter estimates. A list of optimization algorithms for large biological system identification is given elsewhere [16].

#### Step 3: Practical Identifiability Analysis

Once parameters are estimated, practical identifiability analysis is done by constructing a confidence interval of the parameter estimates. In the case of the least-squares/maximum likelihood estimation algorithm, the variance-covariance matrix is estimated by inverting the Fisher Information Matrix [9]. The result of the practical identifiability analysis will give the list of parameters that are not precisely estimated with the provided data and the model structure. When one finds that there is a subset of parameters that are not practically identifiable, the traditional framework recommends conducting experiments that are informative with respect to the practically unidentifiable parameters [14], which is not true in all the cases, because, even with a highly informative data there can practically unidentifiable parameters due to the nature of the model structure/parameter space [15].

#### Step 4: Sensitivity Analysis

Once we obtain the parameter estimates, a local sensitivity analysis is carried out to find sensitive and insensitive parameters in a few cases. The sensitivity analysis result is used to eliminate insensitive/unimportant parameters, which will eventually reduce model complexity.

#### Step 5: Sloppiness Analysis

Along with practical identifiability, sometimes sloppiness analysis is done to find large insensitive directions in the parameter space, even though individual parameters are sensitive. There has been a considerable discussion concerning the utility of sloppiness in system identification. Sloppiness analysis in general carried out locally around the optimal parameter estimates. However, it is not very useful once the estimation is done. The exact source of sloppiness could not be identified as the sloppiness measure defined in is a function of both data and model structure [10].

From steps 1-5, we observe that if the true model is in a sloppy region, using the traditional approach of analysing sloppiness post estimation and using the Hessian of the cost function to assess sloppiness will not identify the source of parameter uncertainty accurately. Hence, we propose a new framework extending ideas we previously proposed [15], to accurately identify the source of parameter uncertainty.

### 2.2 Proposed framework for grey-box identification

In this section, we introduce a new framework for grey-box identification that accurately detects the source of parameter uncertainty in non-linear predictors. Fig. 2 illustrates the flow of the proposed framework.

**Figure 2:**
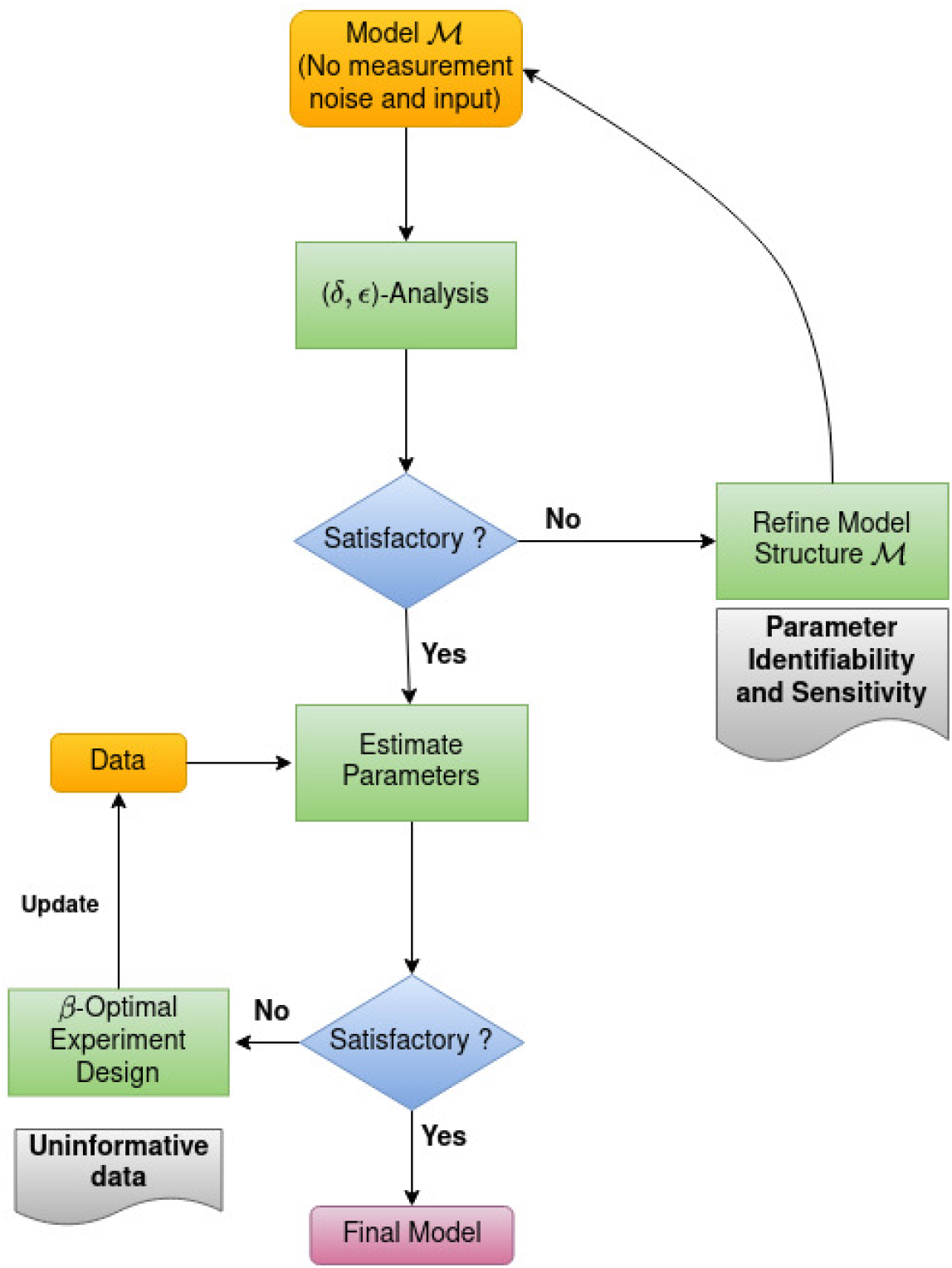
Proposed framework for grey-box identification

#### Step 1: (*δ, ϵ*)-Conditional Sloppiness Analysis

As a first step in conditional sloppiness analysis, the input of the system is switched off, and noise-free data is generated for a sufficiently large duration around a nominal parameter from a viable set of parameters. Then the model sensitivity plot (*δ* vs *ψ*), sloppiness plot (*δ* vs *γ*_*min*_) and (*δ*_*θ*_*i* vs *γ*_*min*_) are plotted for an assumed radius *δ* as proposed in [15]. The volume of the viable parameter space usually dictates the choice of *δ*. The conditional sloppiness analysis is a unified method to detect structural identifiability, sloppiness and parameter sensitivity. If the model is sloppy in stage 1, then a subset of parameters is likely to be practically unidentifiable post-estimation.

#### Stage 2: Parameter Estimation

In this stage, the parameters are estimated with the given experimental condition and practical identifiability analysis is carried out. The source of uncertainty is identified as follows

1. If the model is not conditionally sloppy, and if the parameter estimates are unidentifiable, then the data is not informative.

2. If the model is conditionally sloppy in stage 1, and the parameters are unidentifiable, then first fix the model structure and then, re-estimate the parameters.

From steps 1 and 2, it is clear that the proposed method has a two-fold advantage: (i) it offers a unified framework for detecting identifiability, sloppiness and sensitivity, and (ii) it can detect the source of the parameter uncertainty accurately. In the following section, we demonstrate the working of the proposed method using a numerical example.

## 3 Numerical Example: A simplified model of mRNA Translation

### 3.1 Model Description

In this work, we consider a simplified mRNA transcription model. At *t*_0_, mRNA is released into the cell and gets translated into Green Fluorescent Protein (GFP) at the rate of *p*_3_. Figure 3 shows the mRNA translation and degradation of mRNA and GFP.

**Figure 3:**
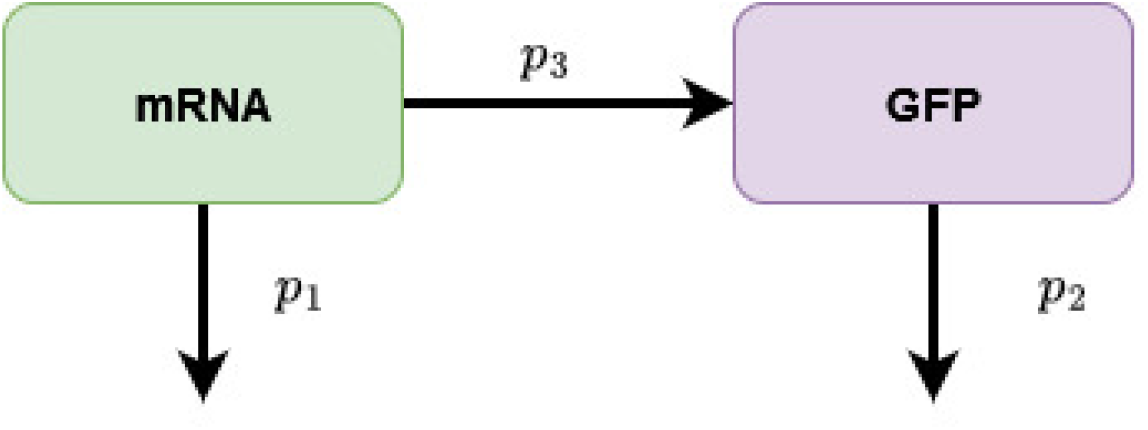
mRNA Transcription Model

The mathematical description as formulated in [8], is given in (1).

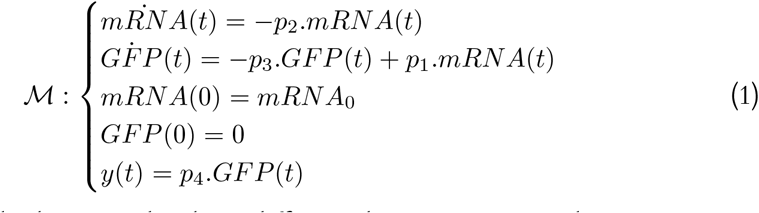

The analytical solution to the above differential equation is given by

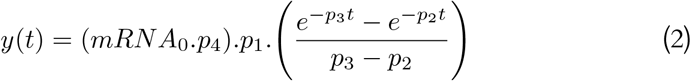

In this study, the initial condition of *mRNA* and *p*_4_ are known and *p*_1_, *p*_2_ and *p*_3_ are the unknowns to be estimated from data.

### 3.2 Data Generation

In this work, we generate data from the differential equation (1) using the parameter values given in Table 1; further, we add Gaussian White Noise to the observation so that the signal-to-noise ratio (SNR) fixed hundred. The model is simulated from *t* = 0 to *t* = 100 seconds with the sampling interval of *t*_*s*_ = 0.5 seconds. Fig. 4 shows the GFP decay along with the measurement noise.

**Table 1:**
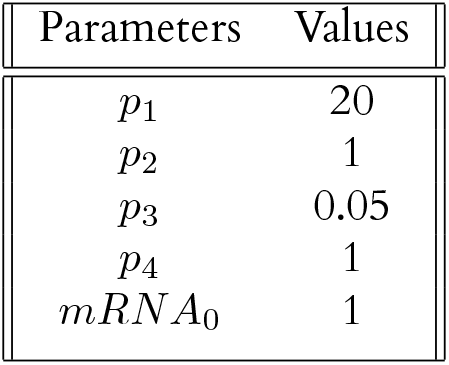
True parameter values.

**Figure 4:**
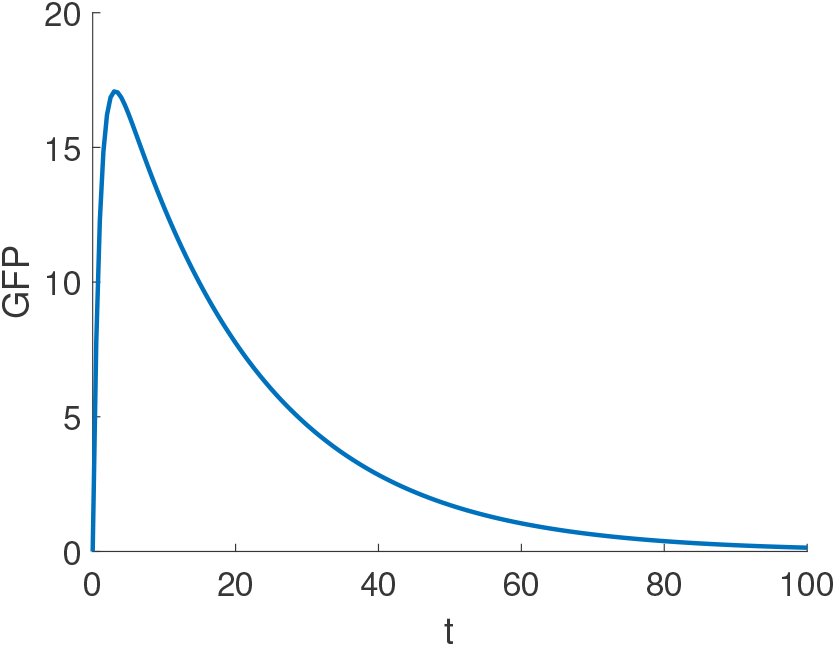
GFP Decay

### 3.3 Structural Identifiability Analysis

In this section, we perform a structural identifiability analysis using GenSSI 2.0 toolbox for MATLAB [17]. A model is structurally unidentifiable if two or more sets of parameters result in identical predictions. There are several methods to detect the structural identifiability of a model structure. This toolbox uses generating series method to identify the identifiability of the parameters. In this case study, we assume that the parameters *p*_4_ and initial condition (*mRNA*_0_) are known. Only parameters in the state equations are considered to be unknown.

From Fig. 5, the algorithm identified *p*_1_ as globally identifiable, and *p*_2_ and *p*_3_ as locally identifiable. It is also clear from (2) that we can swap the values of *p*_2_ and *p*_3_ to get identical predictions. One of the methods for avoiding the local structural unidentifiability in this scenario is to constrain the parameter space, which in this case, it is achieved by taking *p*2 *> p*3.

**Figure 5:**
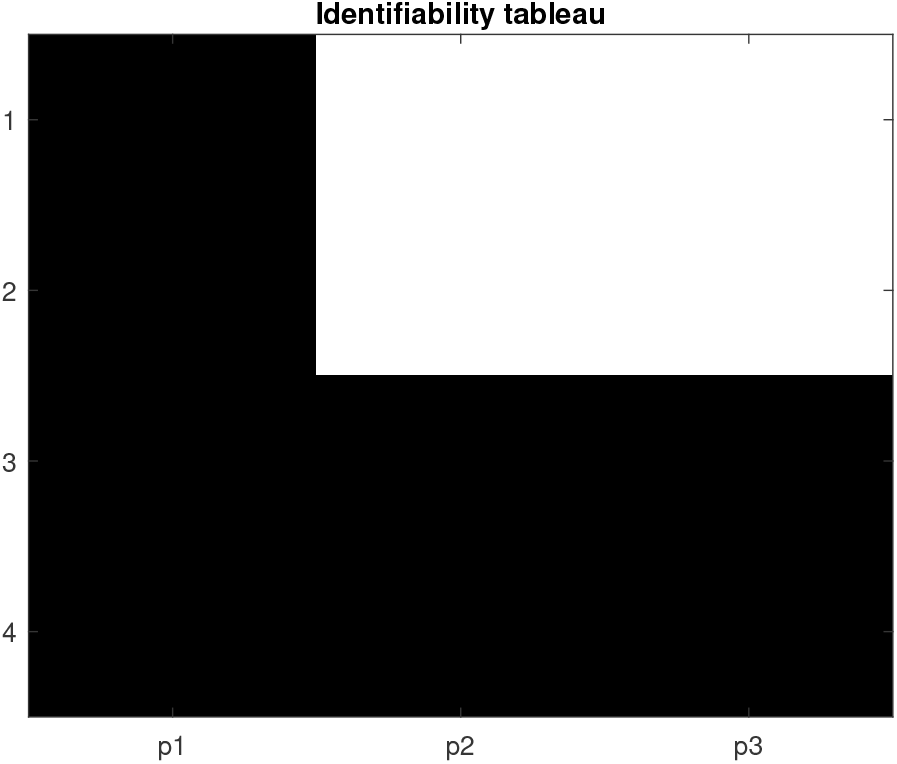
Identifiability Tableau

### 3.4 Parameter and Prediction Uncertainty

In this work, we estimate the parameters using Approximate Bayesian Computation (ABC-MCMC) [11]. Approximate Bayesian Computation is a likelihood free numerical estimation algorithm. All priors are considered to be Gaussian priors.

The posterior distributions and the true parameters are shown in Fig. 6. All the parameters are estimated with low precision despite having a large number of high SNR data points. Also, with reference to the true values in Table 1, the bias in the point estimates for each parameter is significantly high. The primary reason for this is the multi-scale nature of the system.

**Figure 6:**
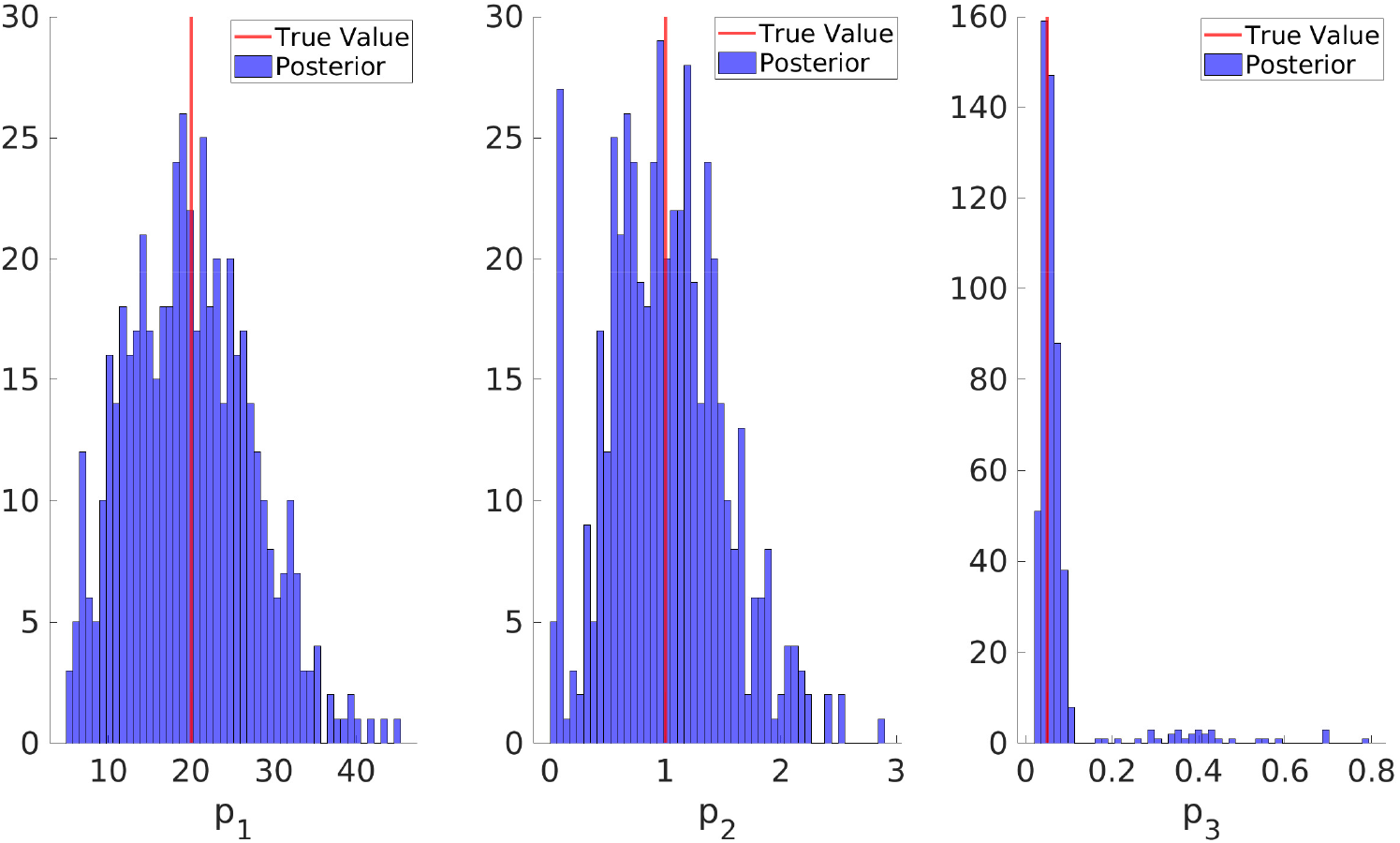
Posterior distributions of parameters

From Fig 7, it can be seen that there is a considerable amount of uncertainty in the transient phase but remains reasonably bounded as the curve approaches a steady state. However, on an average, the predictions are closely tied to the true predictions. We can observe that there exists a subset of predictions that are significantly deviated from the true value because a small set of posterior parameters have fallen into the stiff regimen. This is also a consequence of the presence of sloppiness [10].

**Figure 7:**
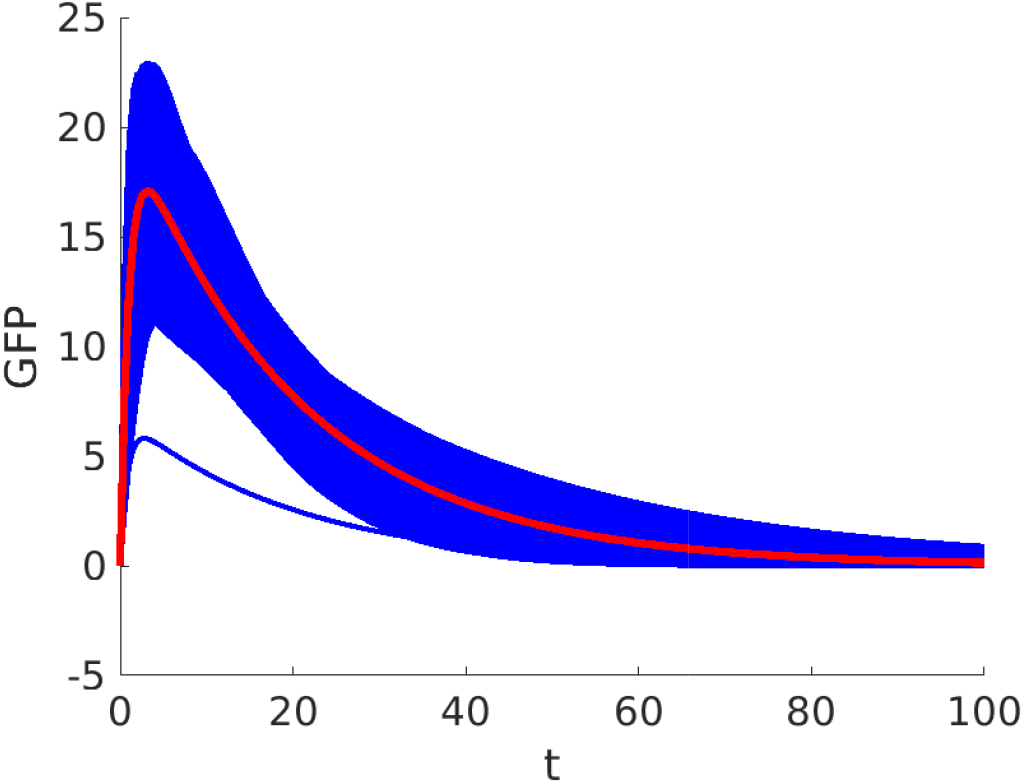
Predictions from all the samples from the posterior distributions.

From Fig. 8, with informative data and locally identifiable model structure, the uncertainty in the predictions and parameter estimates are significant. In this scenario, precisely identifying the source of the errors becomes challenging as the data is informative and the model is identifiable (locally).

**Figure 8:**
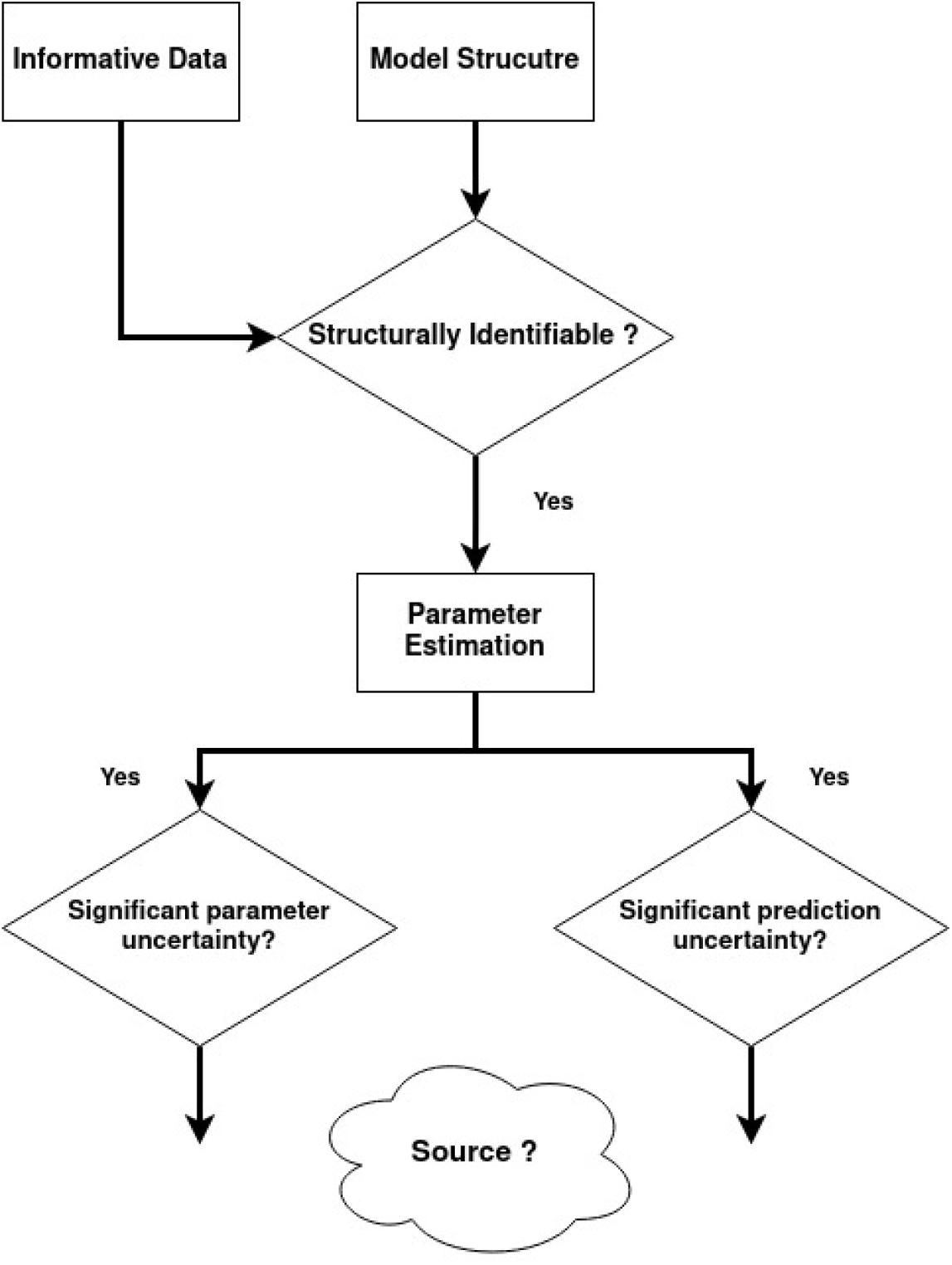
Traditional identification flow for mRNA model

### 3.5 Sensitivity and Sloppiness analysis

In this work, we compute the sensitivity of the output with respect to the parameters. The output parameter sensitivity vector is given by (3). From Eq. (2), we directly compute the partial derivatives of the output with respect to each parameter evaluated at the optimal values obtained in Table 2.

**Table 2:** True parameter values.

| Parameters | Estimates |
| --- | --- |
| $p_1$ | $19.877 (\pm 7.46)$ |
| $p_2$ | $1.001 (\pm 0.502)$ |
| $p_3$ | $0.078 (\pm 0.095)$ |

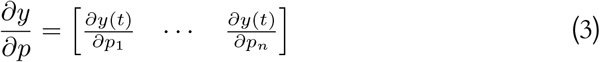

The sensitivity matrix is shown below,

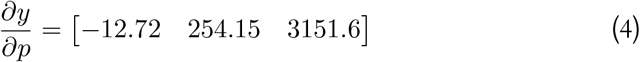

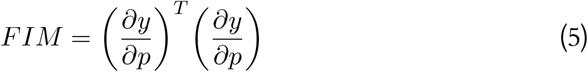

The sensitivity analysis shows that the most sensitive parameter is *p*_3_ followed by *p*_2_ and *p*_1_. However, all the parameters are estimated with low precision. Even with the sensitivity analysis, we cannot find the source. Now we compute the sloppiness analysis by estimating the Fisher Information Matrix from the sensitivity matrix using (5).

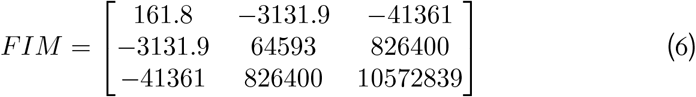

The sloppiness is computed by inverting the condition number of the FIM.

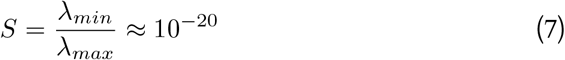

From the sloppiness analysis, we can see that the system is sloppy as *S >* 10^−6^ [10], and that is the reason why we ended up having huge parameter and prediction uncertainty. However, the traditional sloppiness analysis done at the optimal value is both a function of the model and data—hence the ambiguity continues in accurately identifying the source of uncertainty. This is one of the prime reasons why the utility traditional sloppiness measure in system identification is unclear. So, with the traditional identification protocol, in this case, we are unable to accurately identify the source of uncertainty. In order to circumvent this challenge, we now introduce a new framework, where the model structure is tested for the sensitivity of output in the parameter directions prior to the estimation exercise.

### 3.6 Conditional sloppiness analysis

In this section we conduct (*ϵ − δ*) sloppiness analysis as proposed in our previous work [15]. For this analysis, we investigate the model around the true parameter vector. We fix the *δ* = 0.5, the model sensitivity plot and *δ* vs *γ*_*min*_ plots are shown in Figs. 9a and 9b, respectively.

**Figure 9:**
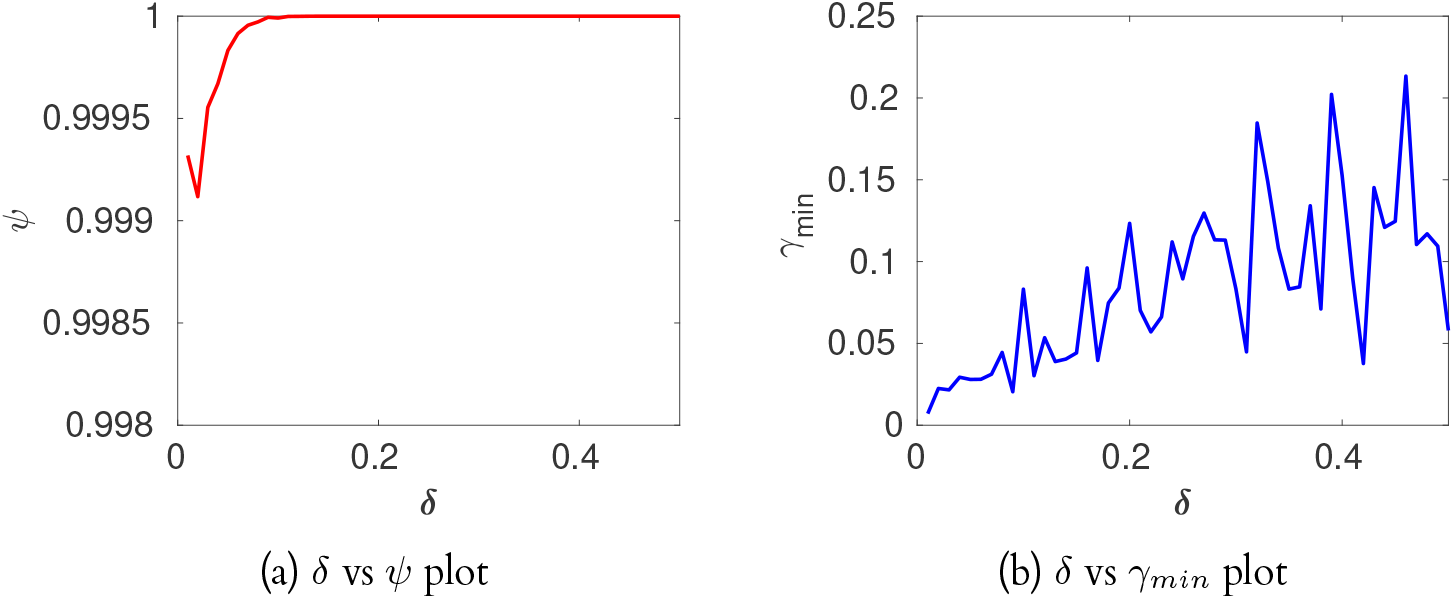
(a) The curve very close to unity value indicating local structural identifiability. (b) The curve has significantly small slope and numerically small *γ*_*min*_ which indicate the sloppiness for the given *δ*

From the above figures, we observe that model has huge anisotropic sensitivity in the observed region; also, from *γ*_*min*_ we can see that the model is conditionally sloppy. This indicates an area in the given *δ* region over which the model predictions are nearly identical, and the region’s geometry has anisotropic sensitivity.

From the plot *δ* versus *γ*_*min*_ in Fig. 10, we observe that the parameter *p*_1_ is highly insensitive for large *δ* up to 0.3, followed by the parameter *p*_2_. This insensitivity is directly reflected in the standard errors of the posterior parameter estimates in Fig. 6. The most sensitive parameter is *p*_3_. For a sufficiently large *δ*, the maximum deviation of *p*_3_ starts to increase exponentially; including such a range of parameters in the posterior distribution of *p*_3_ has resulted in the set of highly uncertain predictions, as seen in Fig. 7.

**Figure 10:**
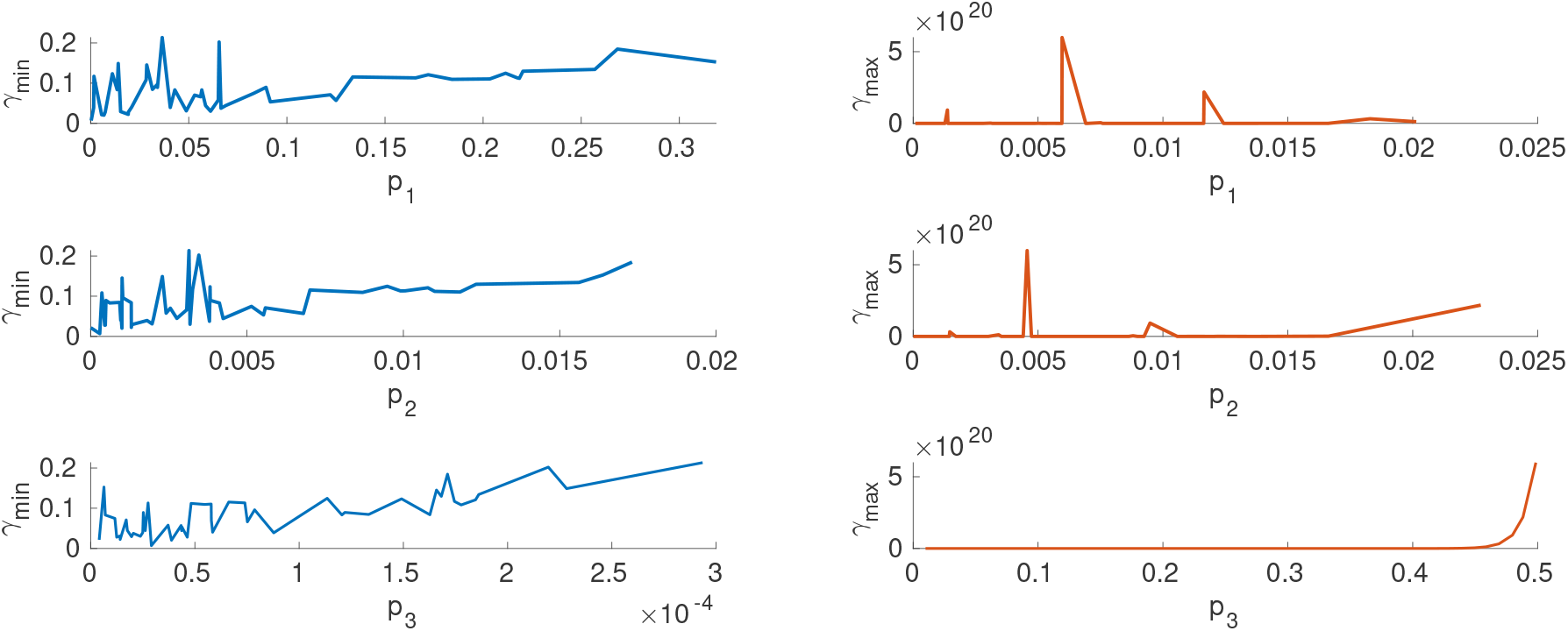
*X*-axis depicts the relative distance of the particular parameter *θ*_*i*_ from its reference value 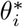. The *Y* -axis on the left is the minimum sum-square deviation and on the right, it is the maximum sum-square deviation from *y*^*^(*t*). Parameterss *p*_1_ and *p*_2_ are more insensitive than *p*_3_.

From the sloppiness analysis, it becomes evident that there is a large region around the true parameter over which the predictions are nearly identical; in other words, even under ideal experimental conditions, the region around the true parameters is sloppy. In this case, the experimental conditions are sampling size and interval, which are sufficiently informative.We also identified insensitive parameters (*p*_1_ and *p*_2_) that are likely to be practically unidentifiable correctly. However, the sensitive parameter *p*_3_ is also practically unidentifiable. This is also one of the classic outcomes of parameter estimation in sloppy models. When the parameter space is highly anisotropic, sometimes during estimation, a small portion of posterior distribution fall into the stiff direction, significantly affecting the prediction uncertainty [10]. Hence, we have accurately identified the model as the source of parameter uncertainty in this case. With the traditional framework, the source of the uncertainty, as demonstrated in the previous section, will remain ambiguous. In this Modeling scenario, one of the ways to circumvent this issue is to choose a tight prior (in the case of Bayesian) or sensitive range (MLE/LS) for the parameters guided by sloppiness analysis during parameter estimation.

## 4 Discussion

Nonlinear system identification poses numerous challenges at each system identification stage; one of the major challenges encountered in grey-box identification is identifying the source of parameter uncertainty. The traditional identification framework might take several iterations to determine the exact source of the problem and, most of the time, results in an ambiguous situation where it becomes nearly impossible to resolve whether data are uninformative or the nature of the model structure is the source of the problem. To circumvent this problem, in this work, we proposed a new framework that includes a unified method to identify sloppiness, identifiability, and sensitivity of the parameters to identify the issues in the model structure. Given a model structure, *M* and data set *z*, structural identifiability analysis and sloppiness analysis are performed around a nominal parameter vector in the viable parameter space as a first step. For this purpose, the observations are generated with no measurement noise, the input is turned off, and the data is collected for a long duration to generate informative data. This step identifies the presence of flat regions in the parameter space and the individual parameter sensitivity. The model structure is then modified if necessary. Once the model structure passes the initial assessment, the parameters are estimated with the given experimental data. If the prediction and parameter uncertainties are not satisfactory, then the source of uncertainty is the data; new experiments can be conducted to maximise the information in the data set. We demonstrated the working of the proposed method with a numerical example where we consider an mRNA transnational network which is multi-scale in nature. This model’s parameter space has sloppy regions, resulting in significant parameter uncertainty even under ideal time series data. In this particular example, sloppiness has a significant impact on the prediction uncertainty because the estimated posterior distribution has a significant overlap with the stiff parameter direction. The proposed framework accurately identified the model as the source of uncertainty. Thus, we believe that the proposed framework has vast potential in grey-box identification of biological processes.

